## Supplemental Data 1 for "Systematically optimized BCMA/CS1 bispecific CAR-T cells robustly control heterogeneous multiple myeloma"

**SUPPLEMENTARY MATERIAL**

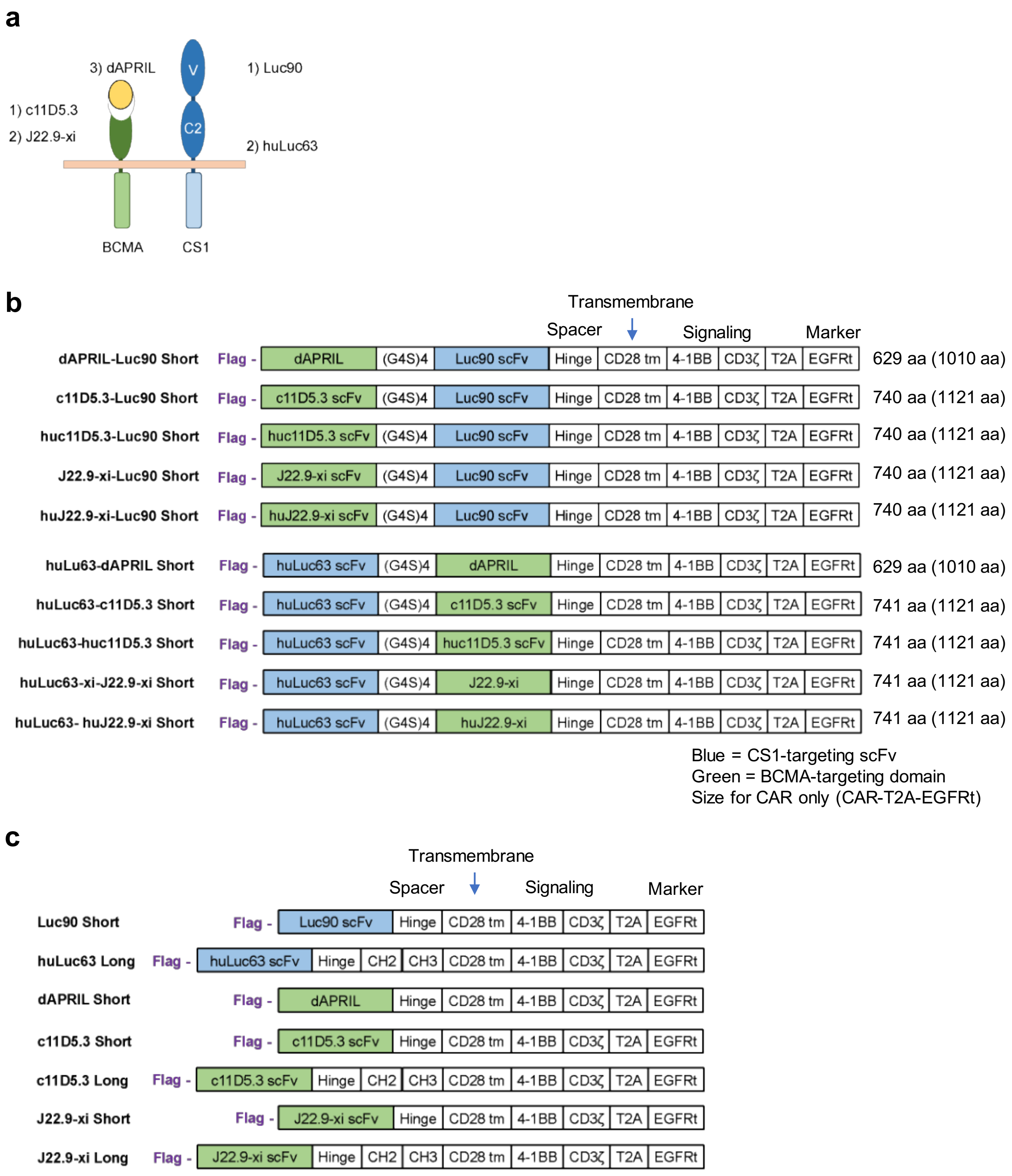


**Supplementary Fig. 1. Rational design of BCMA/CS1 OR-gate CARs and single-input CARs. (a)** OR-gate CARs were constructed based on the binding-epitope location of CS1-specific scFvs. **(b)** Schematic of single-chain bispecific CARs and **(c)** single-input CARs. CARs were tagged with an N-terminal FLAG tag and fused via a self-cleaving T2A peptide to truncated epidermal growth factor receptor (EGFRt), which serves as a transduction marker.

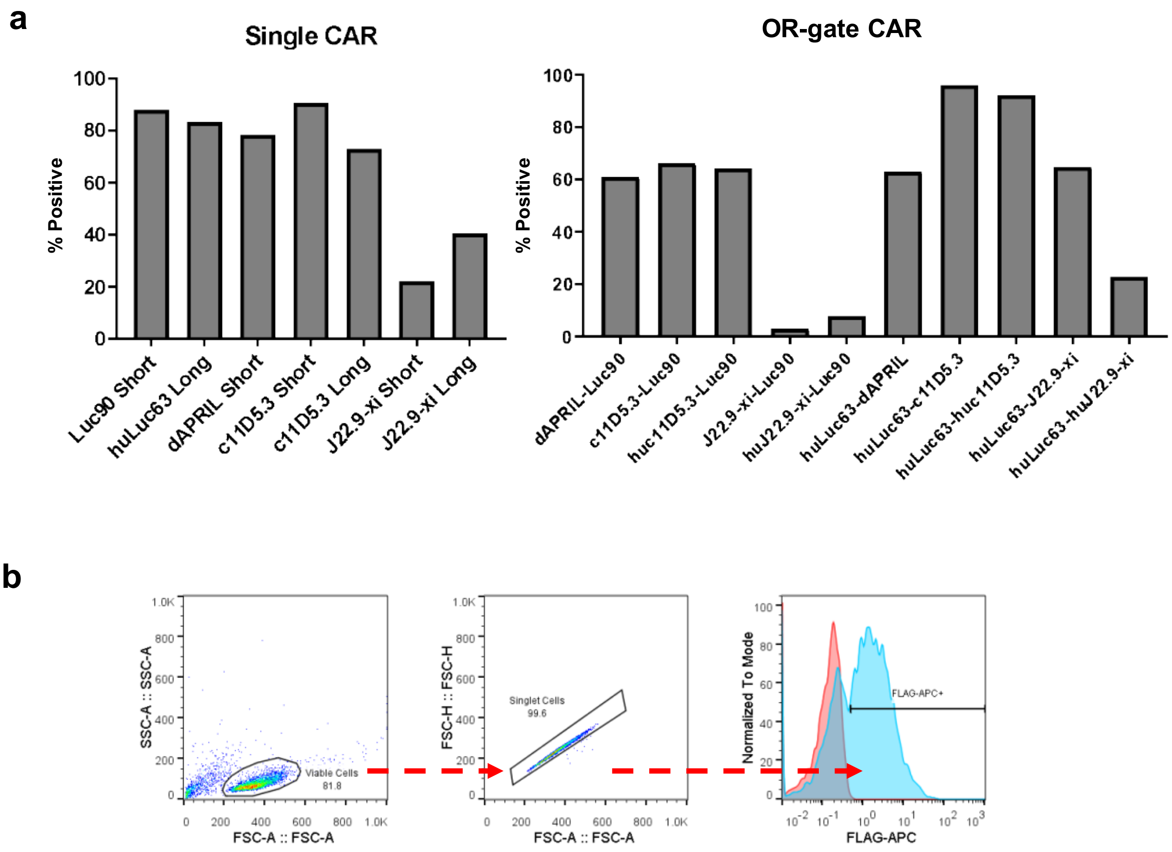


**Supplementary Fig. 2. Bispecific CARs present on primary T-cell surface with varying efficiencies. (a)** FLAG-tagged CAR expression levels were quantified by surface antibody staining followed by flow cytometry. Data shown are representative of results from 3 independent experiments performed with cells from 3 different healthy donors. (**b**) Flow gating strategy for data shown in (a).

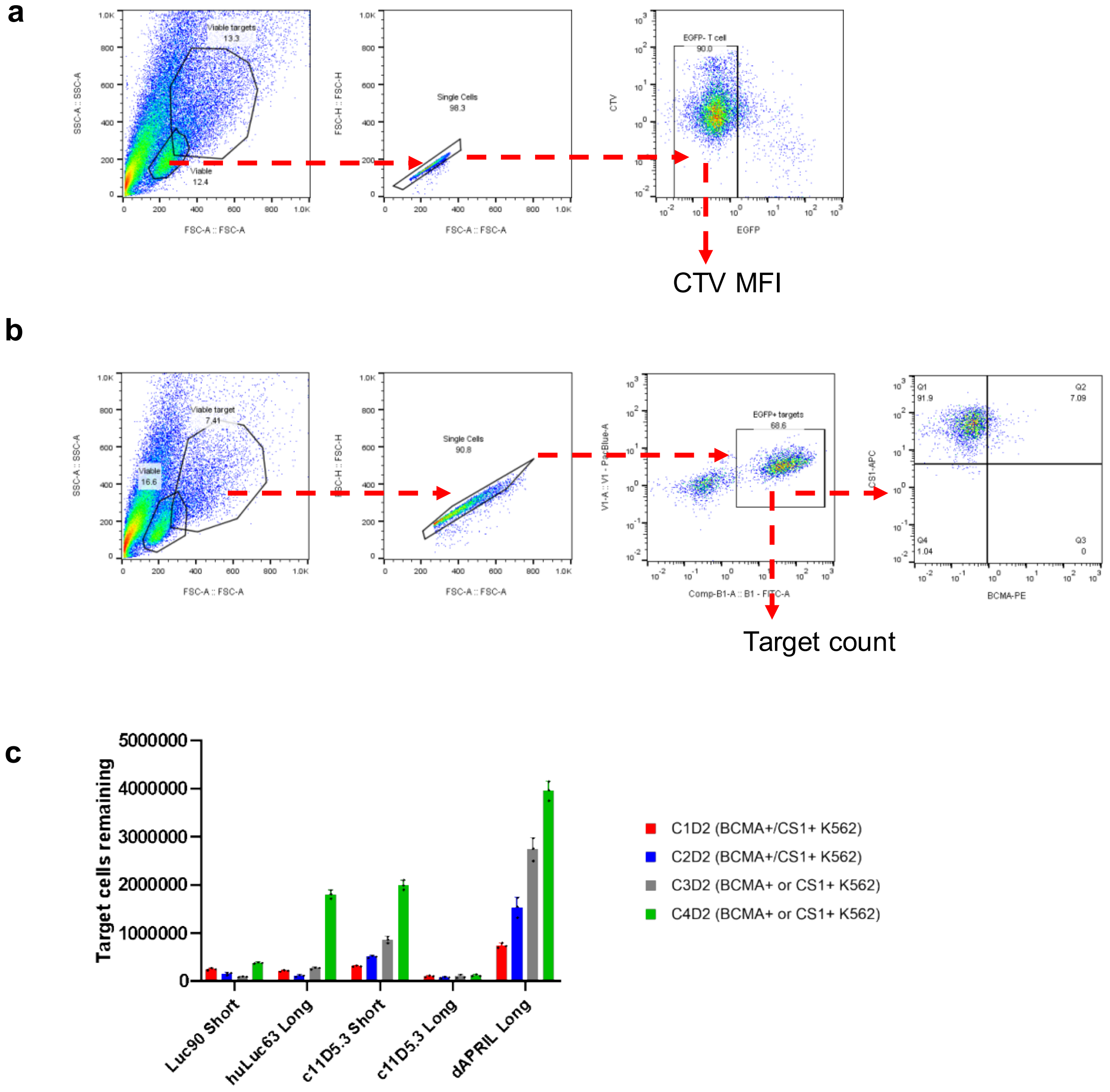


**Supplementary Fig. 3. C11D5.3 scFv shows superior function when separated from the cell membrane by a long extracellular spacer in the CAR context. (a)** Gating strategy for proliferation assay shown in Fig 2b. (**b**) Gating strategy for repeated challenge assay results shown in Fig 2c,d and SFig. 3c. (**c**) Cytotoxicity of single-input CAR-T cells upon repeated antigen challenge. $CD8^+$ CAR-T cells were coincubated with target cells at a 1:1 E:T ratio and re-challenged every 2 days with fresh target cells. BCMA+/CS1+ K562 cells were used as targets for all CAR-T cell samples in challenges 1 and 2. For challenge 3 and 4, single-input CAR-T cells were co-incubated with corresponding antigen-expressing targets (i.e., BCMA CAR-T cells were challenged with BCMA+ K562 cells; CS1 CAR-T cells were challenged with CS1+ K562 cells). Viable target-cell count was quantified by flow cytometry 2 days after each target-cell addition. ‘C#’ denotes the challenge number and ‘D#’ denotes the number of days post challenge. Values shown are the means of technical triplicate samples with error bars indicating + 1 standard deviation (SD).

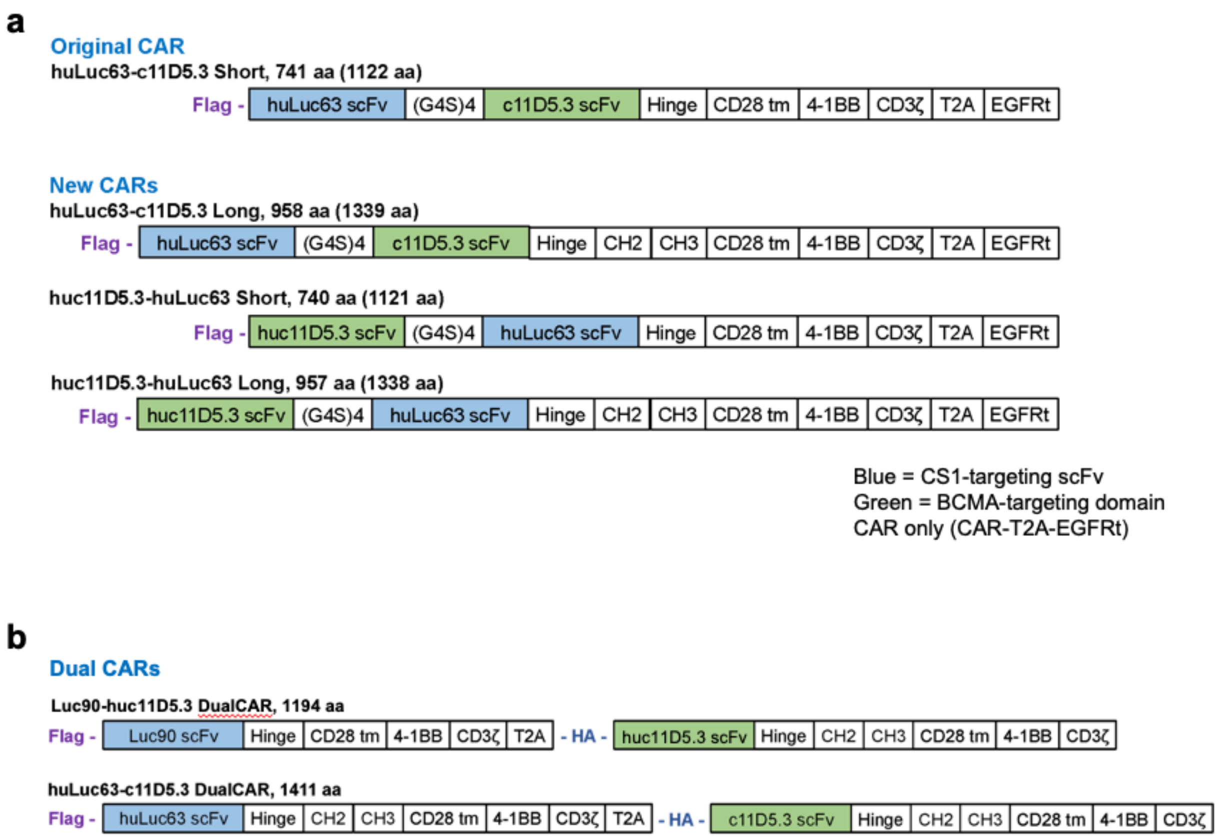


**Supplementary Fig. 4. Schematic of additional OR-gate CAR and DualCAR constructs. (a)** The huLuc63-c11D5.3 Short CAR was modified in an attempt to improve BCMA targeting by replacing the short spacer with a long spacer and/or placing the c11D5.3 scFv in the membrane-distal position. **(b)** DualCAR constructs were constructed where the CS1 CAR was N-terminally tagged with a FLAG tag while the BCMA CAR was N-terminally tagged with a HA tag. The top-performing single-input BCMA CAR identified from previous assays was selected for DualCAR comparison.

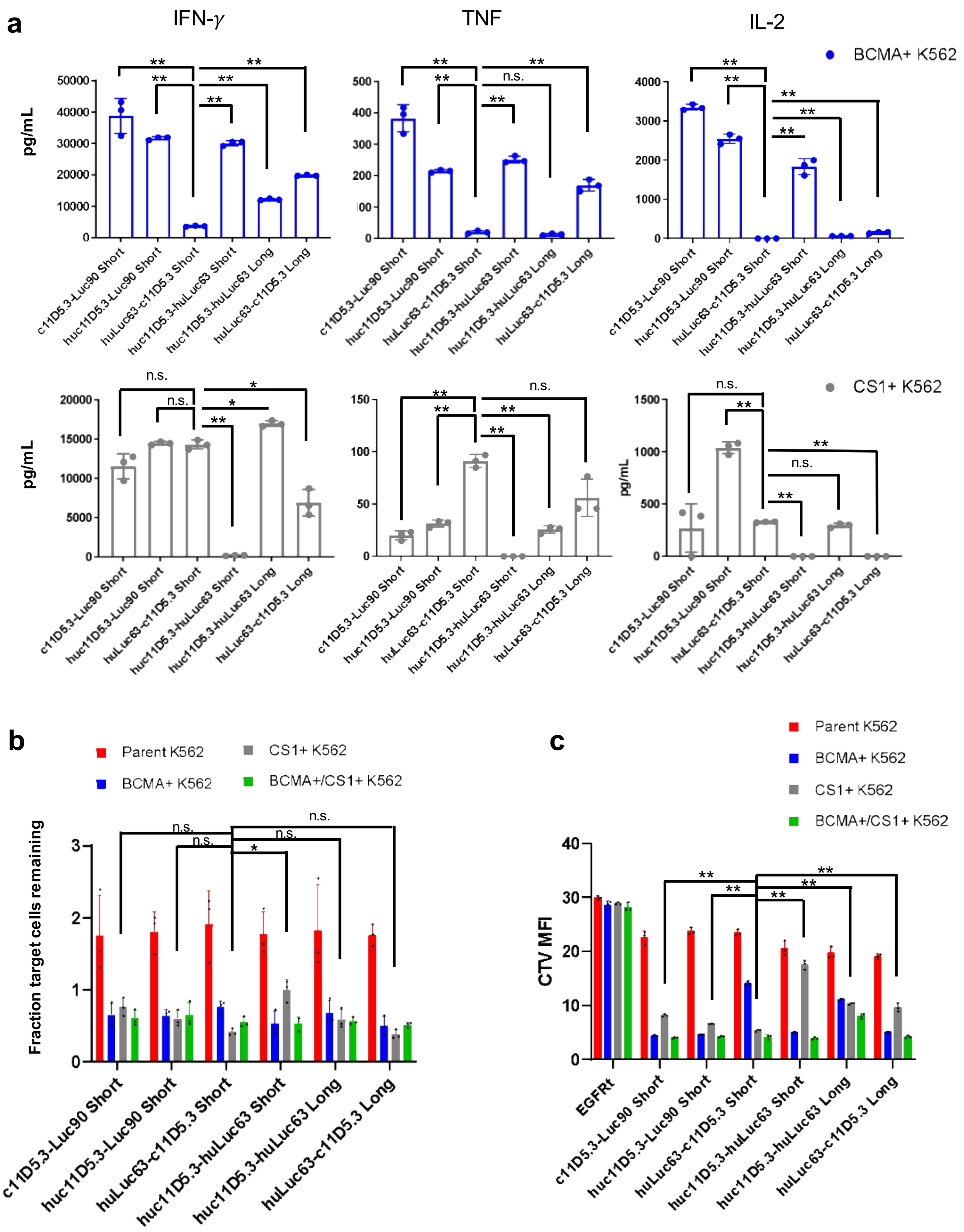


**Supplementary Fig. 5. Modified huluc63-c11D5.3 BCMA/CS1 CARs exhibit improved BCMA-targeting but remains inferior to top-performing candidates. (a)** Cytokine production, **(b)** target-cell lysis, and **(c)** proliferation by CD8$^+$ T cells expressing the modified CAR panel following 24-hour coincubation with BCMA$^+$ or CS1$^+$ K562 cells. Values shown are the means of technical triplicate samples with error bars indicating + 1 SD. *P* values were calculated by unpaired two-tailed Student's $t$-test; n.s. not statistically significant ($p > 0.05$); * $p < 0.05$; ** $p < 0.01$, with Bonferroni correction for multiple comparisons applied.

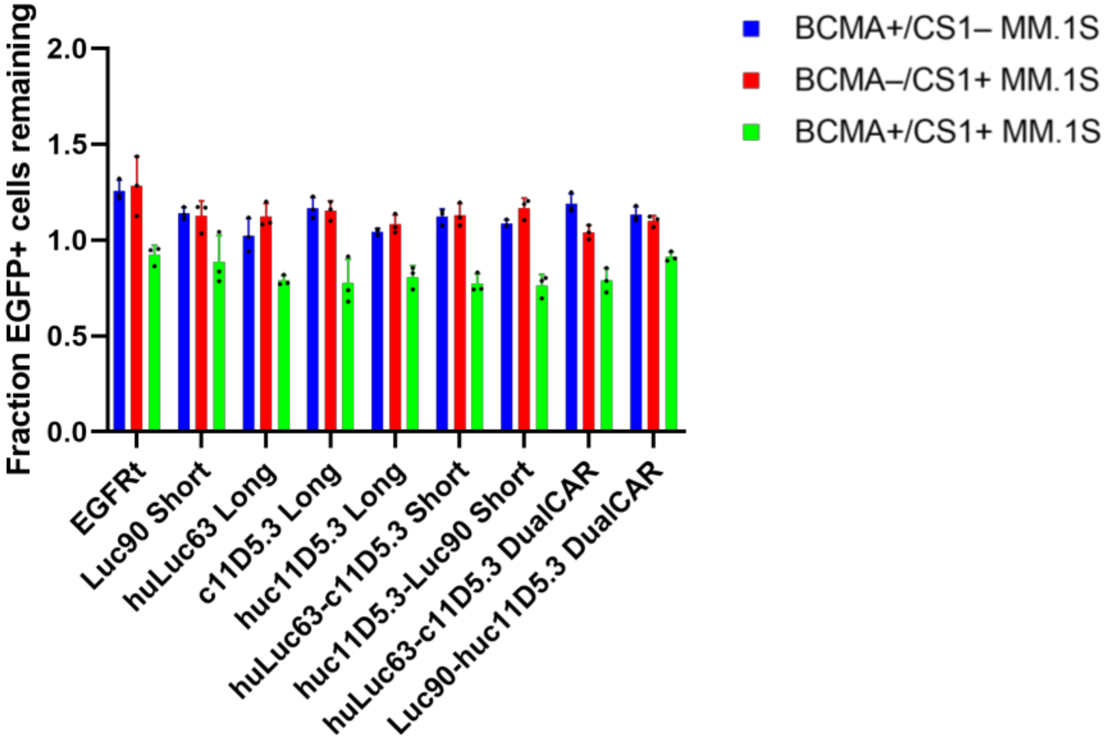


**Supplementary Fig. 6. Low transduction efficiency of DualCAR T cells prevent analysis of target-cell lysis.** T cells transduced with the indicated construct were co-incubated with 5,000 target cells in 96-well plates at a 1:1 E:T ratio, where "effector" was defined as $CAR^+$ T cells. DualCAR T cells exhibited very low transduction efficiency (0.97%–2.56% $DualCAR^+$ across 3 different donors; see Figure 3c). Consequently, a large number of total T cells had to be added to each well to reach the designated E:T ratio while maintaining equal $CAR^+$ (or $EGFRt^+$) cell count as well as equal total T-cell input across samples. Quantification of viable target cell count after a 24-hour co-incubation revealed non-specific killing as the dominant effect observed, resulting in no significant difference between the mock-transduced (EGFRt-only) control compared to any of the CAR-T cell samples. Previous assays had clearly demonstrated functionality of the single-input and single-chain bispecific CARs, thus the lack of relative killing compared to the mock-transduced control observed in this assay is specifically attributable to the high total T-cell input necessitated by the low transduction efficiency of the DualCAR samples. Values shown are the means of technical triplicate samples with error bars indicating + 1 SD.

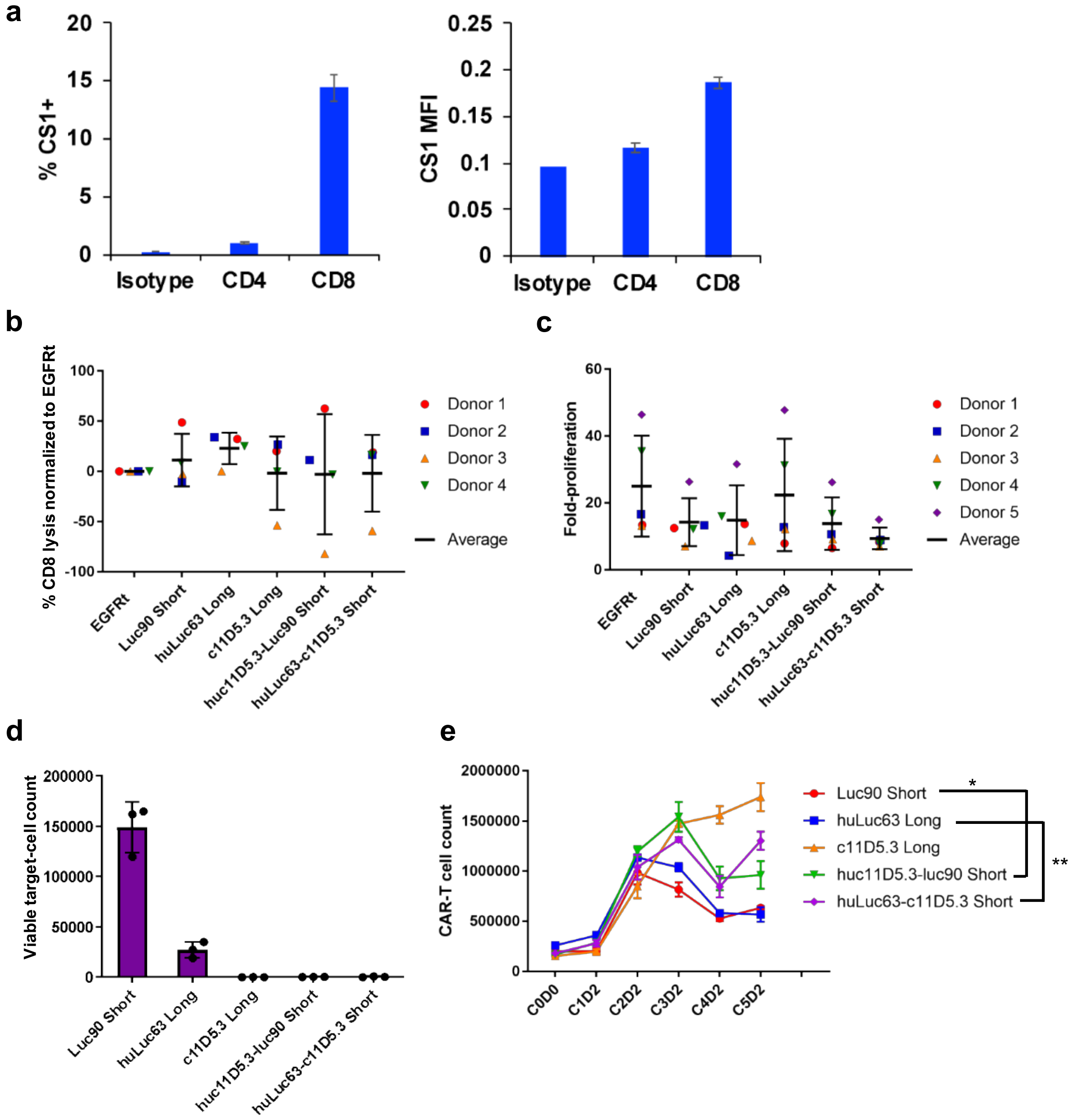


**Supplementary Fig. 7. CS1-specific OR-gate CAR-T cells expand efficiently *ex vivo.*** **(a)** CS1 is expressed at higher levels on CD8$^{+}$ T cells than on CD4$^{+}$ T cells. CS1 expression on CD4$^{+}$ and CD8$^{+}$ T cells from the same donor was quantified by surface antibody staining of CS1 by flow cytometry. **(b)** Fratricide by single-input or bispecific CAR-T cells coincubated with CTV-stained, untransduced (UT) CD8$^{+}$ T cells. Cells were seeded at a 2:1, [CAR-T cells]:[UT-T cell] ratio, and viable CTV-stained UT-T cells were quantified by flow cytometry after a 24-hour coincubation. Percent lysis of UT CD8$^{+}$ T cells was normalized to cells lysed in the EGFRt control group. Values, means, and error bars indicating ± 1 SD from 4 donors are shown. No statistically significant difference in proliferation was observed across the CAR-T cell panel. **(c)** Proliferation of T cells expressing the top-two BCMA/CS1 bispecific CARs or corresponding single-input CARs 10 days post activation by CD3/CD28 Dynabeads. Fold-proliferation is defined as the number of $CAR^+$ T cells on day 10 normalized to the number of T cells on the day of transduction (i.e., 2 days post activation). Values, means, and error bars indicating ± 1 SD from 5 donors are shown. No statistically significant difference in proliferation was observed across the CAR-T cell panel. **(d)** Target-cell lysis by NM single-input and bispecific CARs following repeated antigen challenge. Cells were seeded at a 1:2 E:T ratio. The number of viable $BCMA^+/CS1^+$ MM.1S target cells remaining 2 days after the fifth challenge (i.e., on C5D2) was quantified by flow cytometry. The means of technical triplicate samples are shown with error bars indicating + 1 SD. **(e)** CAR-T cell proliferation over 5 rounds of repeated antigen challenge. Number of CAR-T cells remaining 2 days after each round of antigen challenge was quantified by flow cytometry. Statistics for C5D2 are depicted. *P* values were calculated by unpaired two-tailed Student's *t*-test; n.s. not statistically significant ($p > 0.05$); * $p < 0.05$; ** $p < 0.01$, with Bonferroni correction for multiple comparisons applied.

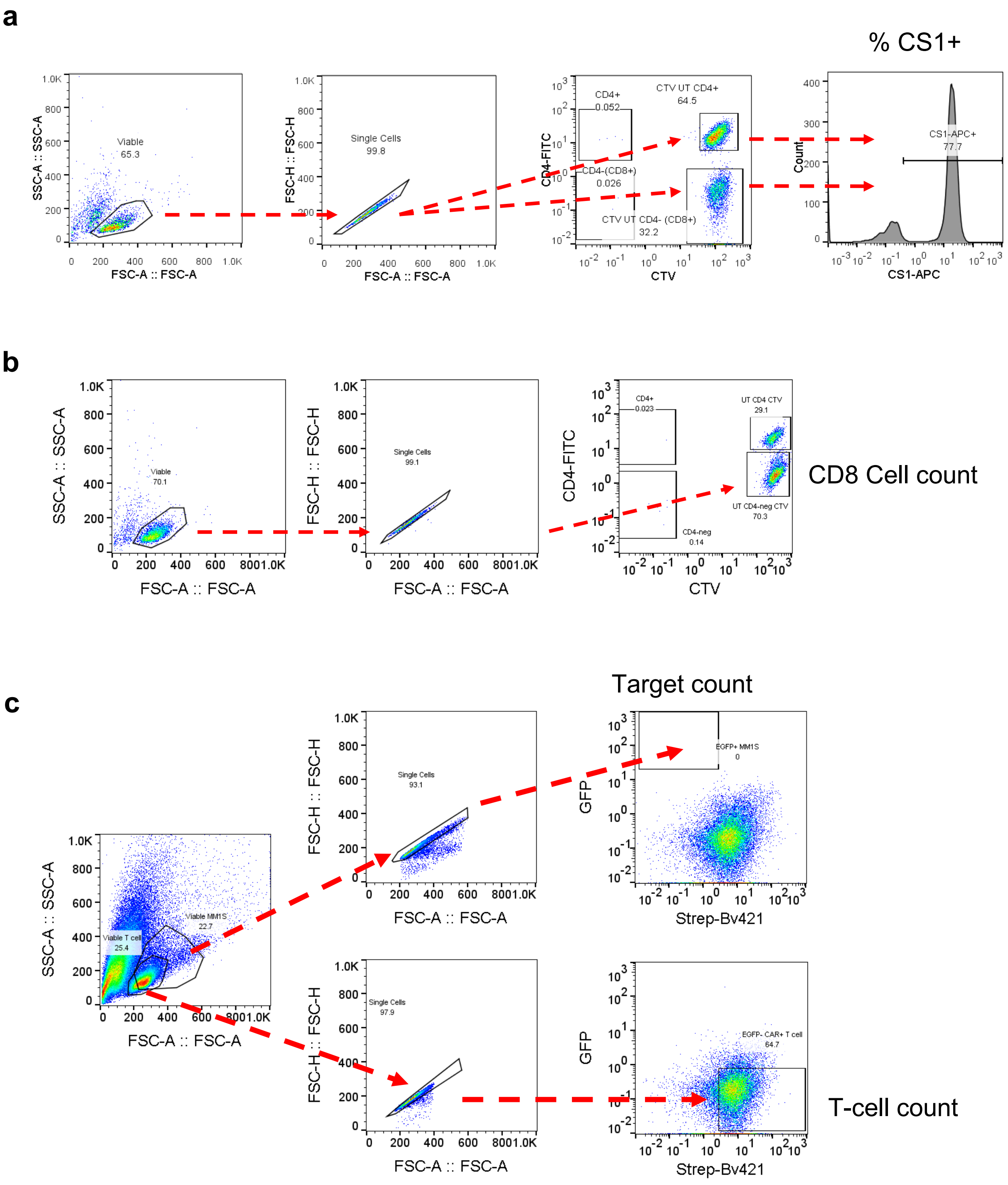


**Supplementary Figure 8. CS1 CAR-T cells exhibit functional defect in repeated antigen challenge**. Gating strategy for data shown in **(a)** Supplementary Fig. 7a, **(b)** Supplementary Fig. 7c, and **(c)** Supplementary Fig. 7d,e.

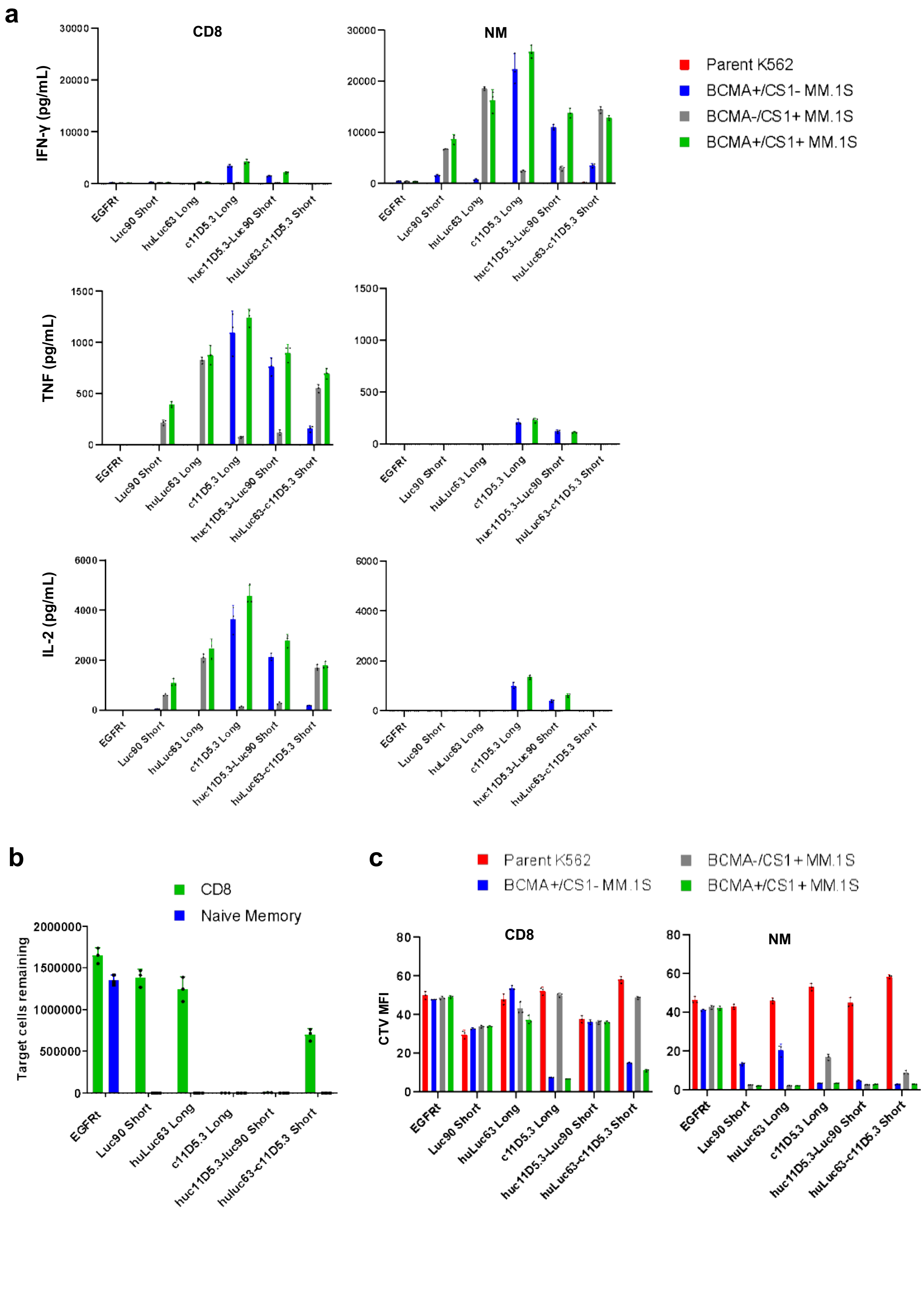
a
CD8
NM
IFN-γ (pg/mL)
TNF (pg/mL)
IL-2 (pg/mL)
Parent K562
BCMA+/CS1- MM.1S
BCMA-/CS1+ MM.1S
BCMA+/CS1+ MM.1S
EGFRt
Luc90 Short
huLuc63 Long
c11D5.3 Long
huc11D5.3-Luc90 Short
huLuc63-c11D5.3 Short
b
CD8
Naive Memory
Target cells remaining
c
CTV MFI
CD8
NM

**Supplementary Fig. 9. Naïve/memory (NM) CAR-T cells exhibit superior *in vitro* anti-tumor killing compared to CD3$^+$ and CD8$^+$ CAR-T cells. (a)** Interferon(IFN)-γ, Tumor Necrosis Factor (TNF), and Interleukin(IL)-2 production of CD8$^+$ or NM single-input and bispecific CAR-T cells following 24-hour coincubation with wildtype (BCMA$^{+/}$CS1$^+$) or CRISPR/Cas9-modified BCMA$^-$/CS1$^+$ and BCMA$^+$/CS1$^-$ MM.1S cells at a 2:1 E:T ratio. **(b)** Target-cell lysis by NM or CD8$^+$ single-input and bispecific CAR-T cells following repeated antigen challenge. Cells were seeded at an E:T ratio of 1:2. Number of remaining BCMA$^+$/CS1$^+$ MM.1S target cells was quantified after 4 rounds of antigen challenge. **(c)** Proliferation of NM or CD8$^+$ single-input and OR-gate CAR-T cells following a 5-day coincubation with target cells. Values shown in bar graphs are the means of technical triplicate samples from the same donor, with error bars indicating + 1 SD.

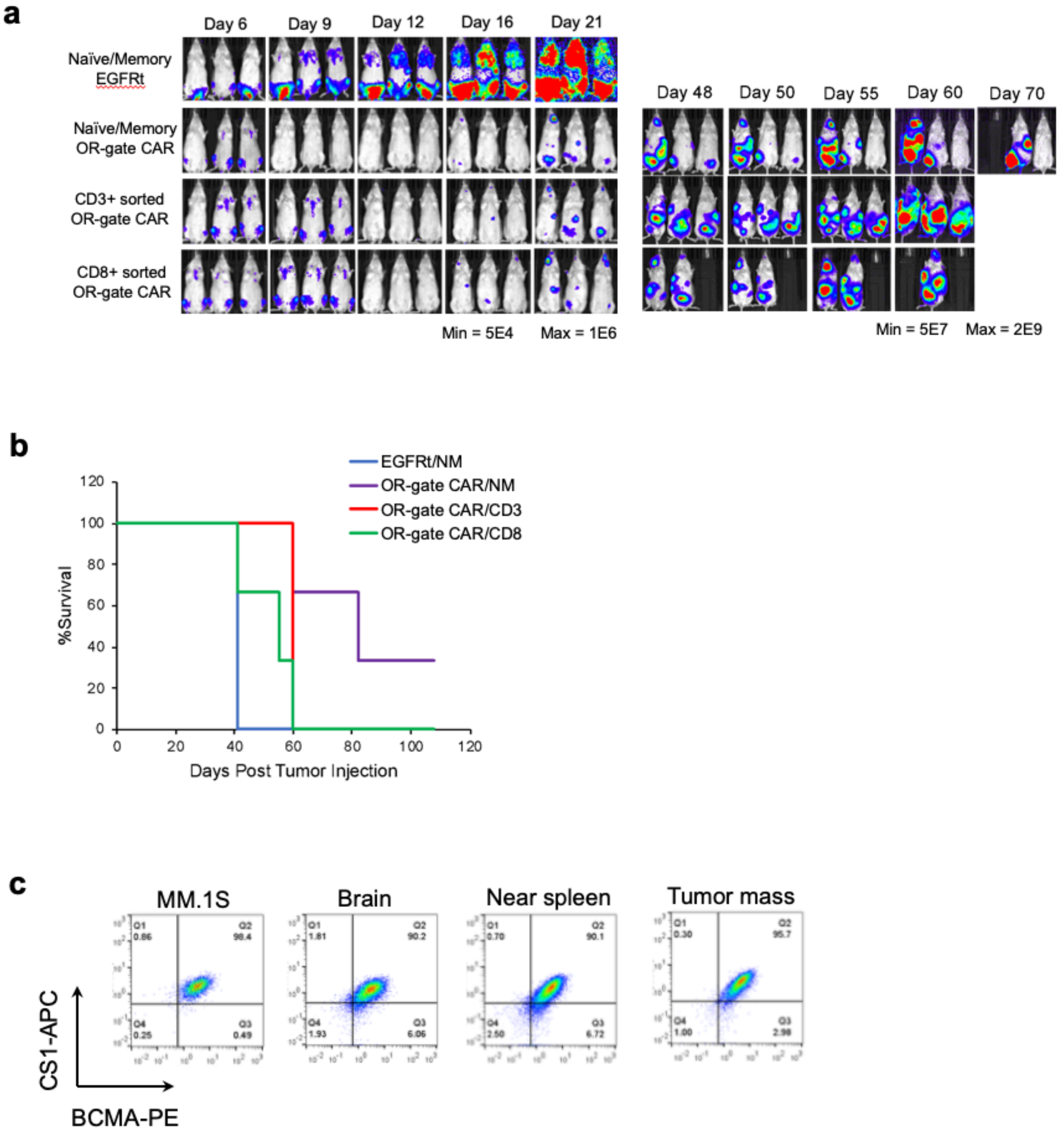


**Supplementary Fig. 10. NM CAR-T cells exhibit superior *in* vivo anti-tumor killing. (a)** Evaluation of NM, CD8$^{+}$, and CD3$^{+}$ huc11D5.3-Luc90 CAR-T cells *in vivo.* NSG mice were engrafted with 2x10$^{6}$ WT MM.1S cells, and 0.5x10$^{6}$ T cells of the indicated subtype and CAR expression were injected 6 days post tumor injection. Tumor progression was tracked by bioluminescence imaging. (**b)** Survival curve of mice shown in (a). The small sample size did not permit detection of statistical significance. However, the last mouse in the NM/OR-gate CAR treatment group had been tumor-free for 53 days and was healthy at the time of study termination on day 108. (**c**) BCMA and CS1 expression on tumor cells isolated from relapsed animals. Data shown are tumors isolated from mouse 1 treated with NM CAR-T cells (sacrificed on Day 60).

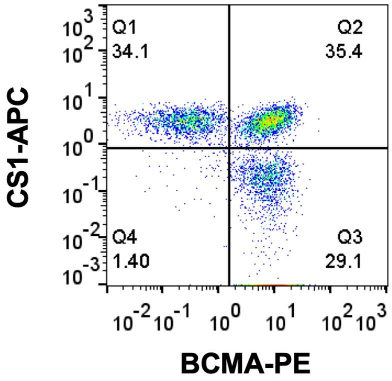


**Supplementary Fig. 11. Mice were injected with BCMA$^+$/CS1$^-$, BCMA$^-$/CS1$^+$, and BCMA$^+$/CS1$^+$ cells mixed at a 1:1:1 ratio.**

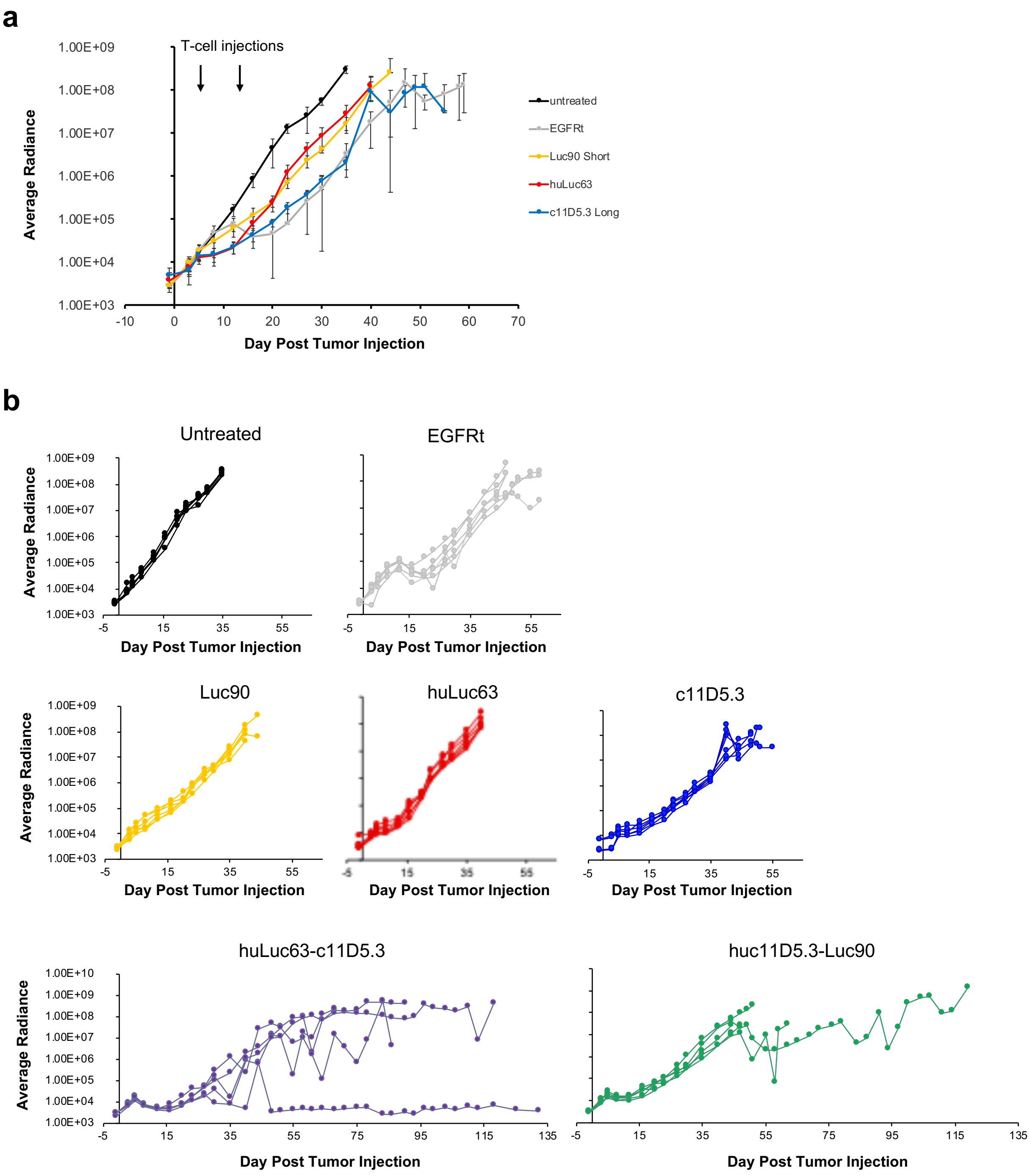


**Supplementary Fig. 12. Tumor progression in CAR-T cell-treated mice. (a)** Average tumor radiance in mice treated with single-input CAR-T cells (n = 6). **(b)** Tumor radiance in individual mice from all treatment groups. Six mice were included in all treatment groups except for huLuc63-c11D5.3, where only 5 were retreated due to limited T-cell availability. One mouse in the huLuc63-c11D5.3 treatment group exhibited complete tumor clearance, and no tumor cells were detected at the time of conclusion of the study.

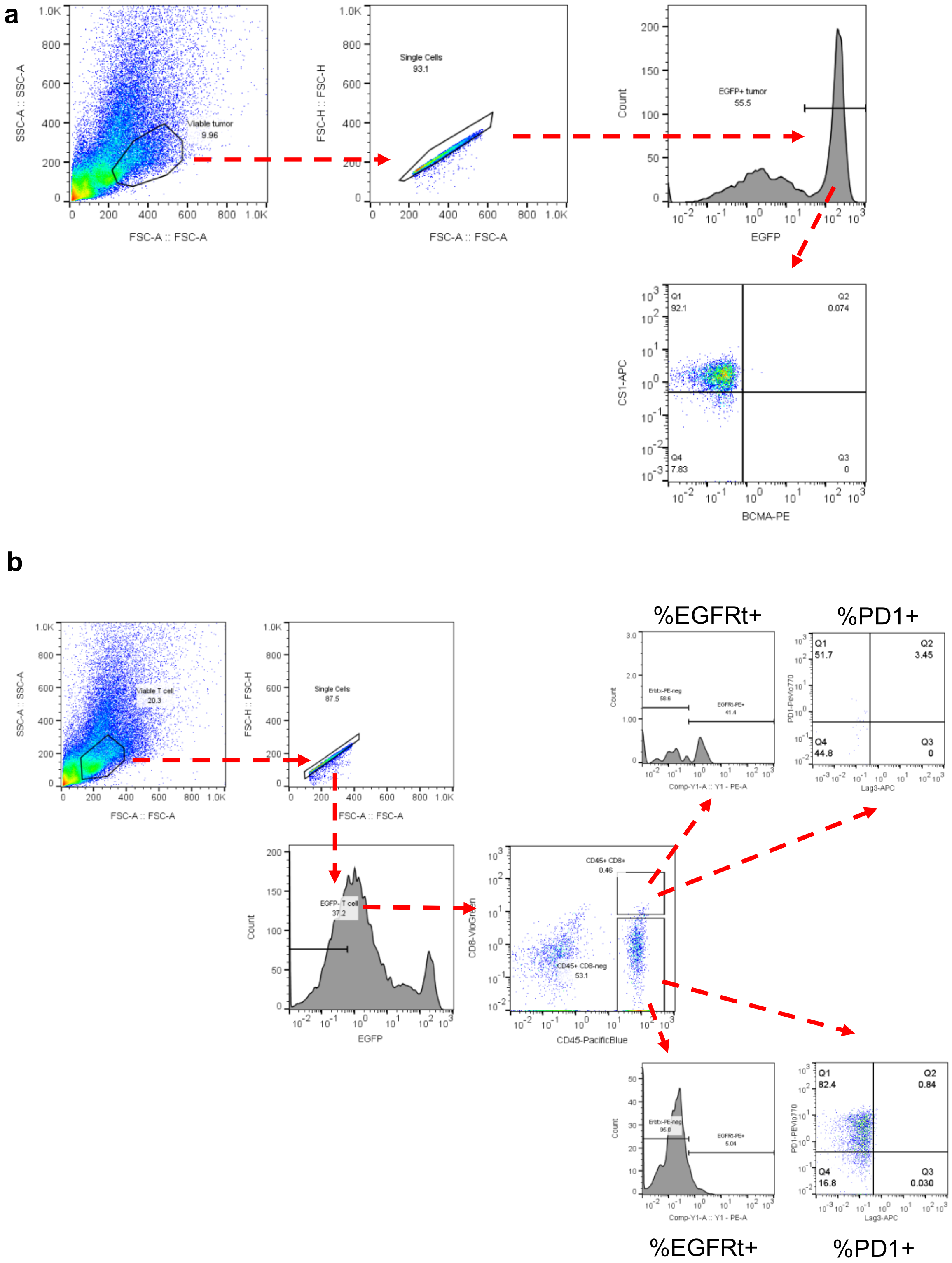


**Supplementary Fig. 13**. **Analysis of persisting T cells and tumor cells in mice treated with OR-gate CARs.** (**a)** Gating strategy to determine phenotype of persisting tumor cells harvested at the time of animal sacrifice. (**b**) Gating strategy to determine phenotype of persisting T cells.

### Mouse #5 Tumor

```
   WT 1    GCTCTTGCTGCATTTGCTCTGGAATTCTTGTAGAGATATTACTTGTCCTTCCAGGCTGTT  60
           ||||||||||||||||||||||||||||||||||||||||||||||||||||||||||||
Tumor 1    GCTCTTGCTGCATTTGCTCTGGAATTCTTGTAGAGATATTACTTGTCCTTCCAGGCTGTT  60

   WT 61   CTTTCTGTAGCTCCCTTGTTTTCTTTTTGTGATCATGTTGCAGATGGCTGGGCAGTGCTC  120
           ||||||||||||||||||||||||||||||||||||||||||||||||||||||||||||
Tumor 61   CTTTCTGTAGCTCCCTTGTTTTCTTTTTGTGATCATGTTGCAGATGGCTGGGCAGTGCTC  120

   WT 121  CCAAAATGAATATTTTGACAGTTTGTTGCATGCTTGCATACCTTGTCAACTTCGATGTTC  180
           ||||||||||||||||||||||||||||||||||||||||||||||||||||||||||||
Tumor 121  CCAAAATGAATATTTTGACAGTTTGTTGCATGCTTGCATACCTTGTCAACTTCGATGTTC  180

   WT 181  TTCTAATACTCCTCCT--CTAACATGTCAGCGTTATTGTAATGCAAGTAAGTAATATTGC  238
           ||||||||||||||||  ||||||||||||||||||||||||||||||||||||||||||
Tumor 181  TTCTAATACTCCTCCTAGCTAACATGTCAGCGTTATTGTAATGCAAGTAAGTAATATTGC  240

   WT 239  TTGAACGATTATTCATTGGTGTGAACTATTCTGTCTATATGGACTGCTTATTCAGAGAAT  298
           ||||||||||||||||||||||||||||||||||||||||||||||||||||||||||||
Tumor 241  TTGAACGATTATTCATTGGTGTGAACTATTCTGTCTATATGGACTGCTTATTCAGAGAAT  300

   WT 299  CAACATAATGGGCATGATGGTGAGTTTTCTTGAATCAAAAAGAGAAAGGAAGCAAGGCAG  358
           ||||||||||||||||||||||||||||||||||||||||||||||||||||||||||||
Tumor 301  CAACATAATGGGCATGATGGTGAGTTTTCTTGAATCAAAAAGAGAAAGGAAGCAAGGCAG  360

   WT 359  TGATT   363
           |||||
Tumor 361  TGATT   365
```

### Mouse #6 Tumor

```
   WT 1    GCTCTTGCTGCATTTGCTCTGGAATTCTTGTAGAGATATTACTTGTCCTTCCAGGCTGTT  60
           ||||||||||||||||||||||||||||||||||||||||||||||||||||||||||||
Tumor 1    GCTCTTGCTGCATTTGCTCTGGAATTCTTGTAGAGATATTACTTGTCCTTCCAGGCTGTT  60

   WT 61   CTTTCTGTAGCTCCCTTGTTTTCTTTTTGTGATCATGTTGCAGATGGCTGGGCAGTGCTC  120
           ||||||||||||||||||||||||||||||||||||||||||||||||||||||||||||
Tumor 61   CTTTCTGTAGCTCCCTTGTTTTCTTTTTGTGATCATGTTGCAGATGGCTGGGCAGTGCTC  120

   WT 121  CCAAAATGAATATTTTGACAGTTTGTTGCATGCTTGCATACCTTGTCAACTTCGATGTTC  180
           ||||||||||||||||||||||||||||||||||||||||||||||||||||||||||||
Tumor 121  CCAAAATGAATATTTTGACAGTTTGTTGCATGCTTGCATACCTTGTCAACTTCGATGTTC  180

   WT 181  TTCTAATACTCCTCCTCTAACATGTCAGCGTTATTGTAATGCAAGTAAGTAATATTGCTT  240
           |||||||            |||||||||||||||||||||||||||||||||||||||||
Tumor 181  TTCTAAT------------ACATGTCAGCGTTATTGTAATGCAAGTAAGTAATATTGCTT  228

   WT 241  GAACGATTATTCATTGGTGTGAACTATTCTGTCTATATGGACTGCTTATTCAGAGAATCA  300
           ||||||||||||||||||||||||||||||||||||||||||||||||||||||||||||
Tumor 229  GAACGATTATTCATTGGTGTGAACTATTCTGTCTATATGGACTGCTTATTCAGAGAATCA  288

   WT 301  ACATAATGGGCATGATGGTGAGTTTTCTTGAATCAAAAAGAGAAAGGAAGCAAGGCAGTG  360
           ||||||||||||||||||||||||||||||||||||||||||||||||||||||||||||
Tumor 289  ACATAATGGGCATGATGGTGAGTTTTCTTGAATCAAAAAGAGAAAGGAAGCAAGGCAGTG  348

   WT 361  ATT   363
           |||
Tumor 349  ATT   351
```

**Supplementary Fig. 14. Residual tumors recovered from animals treated with OR-gate CAR-T cells contained mutations in the CRISPR-targeted BCMA site, indicating origin from the engineered BCMA–/CS1+ MM.1S cell line.** Two tumor samples (87.2% and 88.9% BCMA$^-$/CS1$^+$, respectively) recovered from two different animals in the huLuc63-c11D5.3 Short CAR-T cell-treated group shown in Fig. 5 were expanded for 21 days to obtain sufficient numbers for analysis by amplicon sequencing. The most dominant BCMA sequence in each tumor sample (accounting for 99.5% and 99.7% of all reads in each sample, respectively) is shown in alignment with the WT BCMA sequence within the sequenced amplicon. Green and yellow highlights mark the PAM and guide RNA sequences used in CRISPR/Cas9-mediated gene editing in order to generate the $BCMA^{-}/CS1^{+}$ MM.1S line used in the animal study.

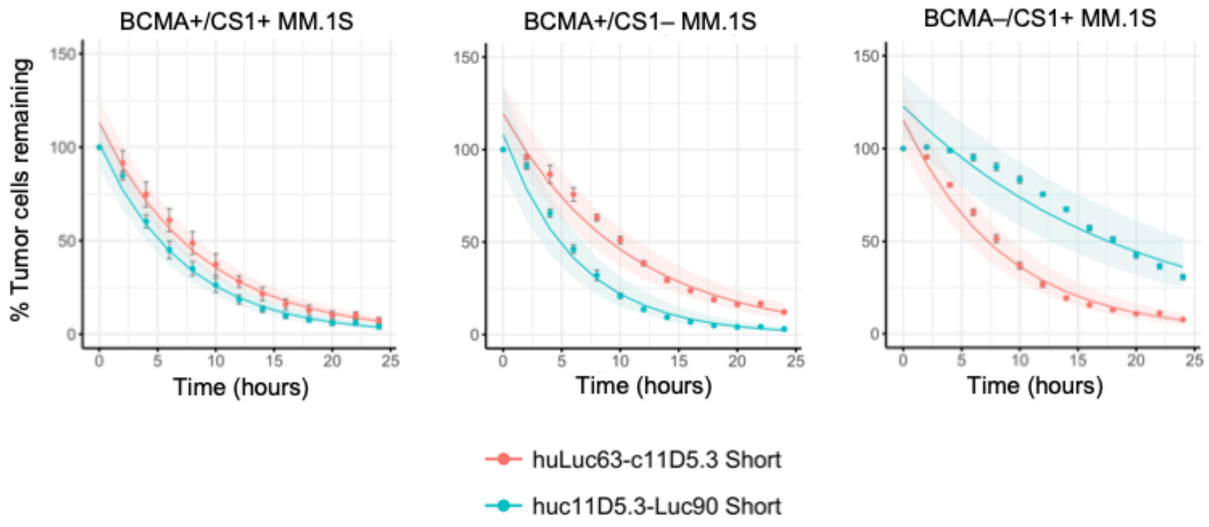


| CAR-T Cell | Kill Rate Constants ± SE (x10$^{-2}$/hr) | | |
|---|---|---|---|
| | BCMA+/CS1+ | BCMA+/CS1– | BCMA–/CS1+ |
| EGFRt | 1.36 ± 0.04 | 1.04 ± 0.05 | 1.12 ± 0.05 |
| Luc90 | 8.40 ± 0.41 | 2.07 ± 0.30 | 8.30 ± 0.34 |
| huLuc63 | 13.76 ± 0.36 | 1.35 ± 0.18 | 13.12 ± 0.47 |
| c11D5.3 | 11.33 ± 0.29 | 13.09 ± 0.57 | 1.56 ± 0.20 |
| huc11D5.3-Luc90 | 13.76 ± 0.40 | 15.98 ± 0.66 | 5.10 ± 0.43 |
| huLuc63-c11D5.3 | 11.56 ± 0.33 | 9.55 ± 0.40 | 11.55 ± 0.47 |

**Supplementary Fig. 15. Kill rate constants of top-performing OR-gate CARs.** Killing of MM.1S tumor cells by OR-gate CAR-T cells was evaluated over a period of 24 hours by IncuCyte. Kill rate constant with standard error (SE) was determined by fitting the data to a log-linear model in R. The mean % tumor cells remaining across time from technical triplicate samples are shown, with the model overlaid and error bars representing ± 1 SD. Shading indicates the 95% confidence interval of the model's fit.

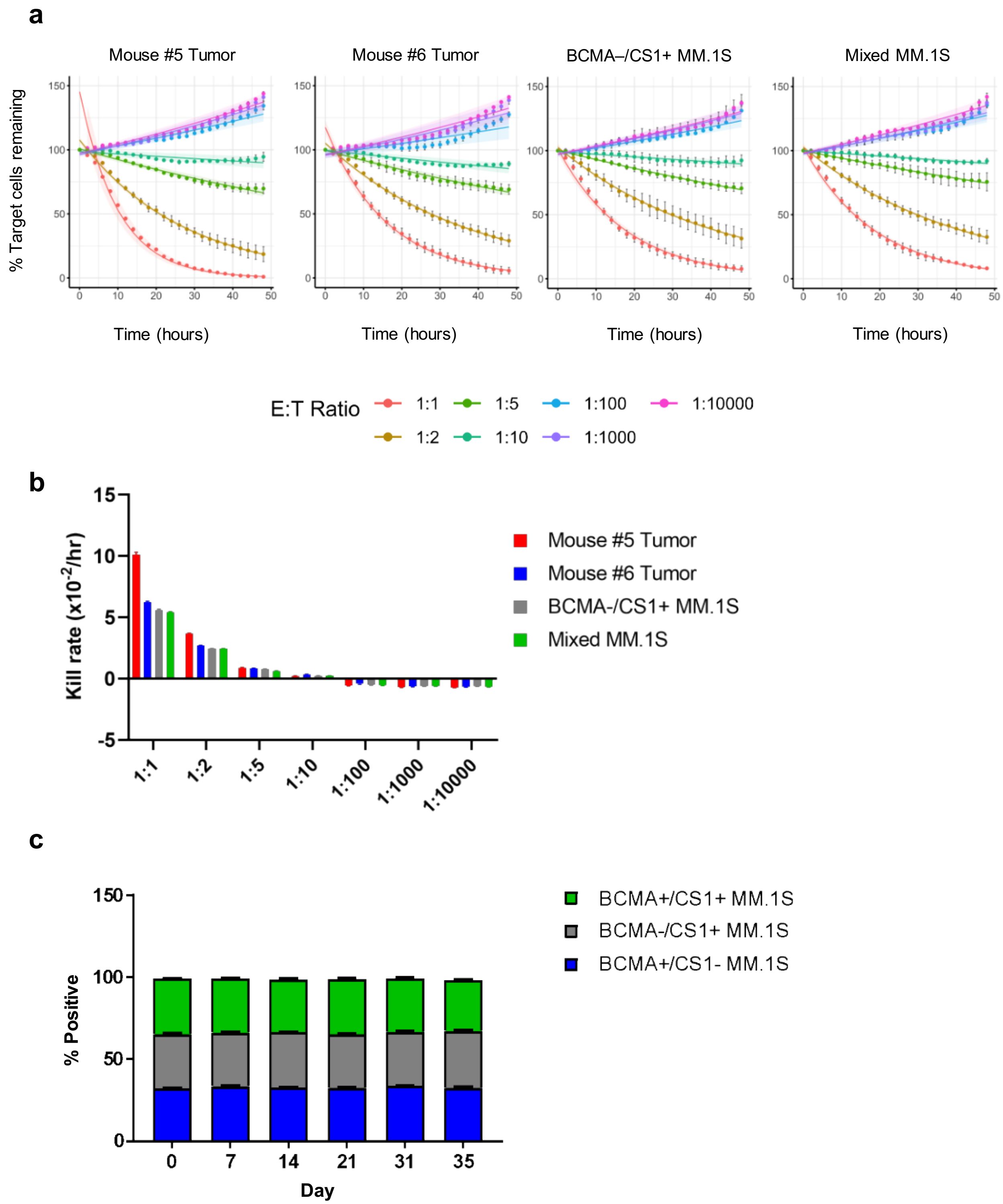


**Supplementary Fig. 16. Recovered tumor cells remain targetable by OR-gate CAR-T cells, and BCMA and CS1 expression pattern does not affect *in vitro* MM.1S growth rate. (a)** Tumor cells recovered from OR-gate CAR-T cell-treated mice remain susceptible to OR-gate CAR-T cell-mediated killing. MM.1S tumor cells recovered from mice #5 and #6 in the huLuc63-c11D5.3 Short CAR-T cell-treated group shown in Figure 4 were sorted to purity based on EGFP expression, expanded for 21 days to obtain sufficient cell numbers for analysis, and subjected to coincubation with huLuc63-c11D5.3 Short CAR-T cells at multiple E:T ratios. $BCMA^{-}/CS1^{+}$ MM.1S cells and a mixture of 1:1:1 of $BCMA^{+}/CS1^{+}$, $BCMA^{-}/CS1^{+}$, and $BCMA^{+}/CS1^{-}$ MM.1S cells were included as controls. Results indicate the tumor cells remain vulnerable to killing by the OR-gate CAR-T cells. The mean % tumor cells remaining across time from technical triplicate samples are shown as described in Supplementary Fig. 13. **(b)** Kill rates (± SE) of the data shown in (A) were determined by fitting the data to a log-linear model in R. **(c)** $BCMA^{+}/CS1^{-}$, $BCMA^{-}/CS1^{+}$, and $BCMA^{+}/CS1^{+}$ MM.1S cells were combined at equal ratios (1:1:1) and co-cultured for 35 days. The % of each cell type quantified throughout the co-culture period showed equal growth rates for the three MM.1S cell lines. Values shown are the means of technical triplicates with error bars indicating + 1 SD.

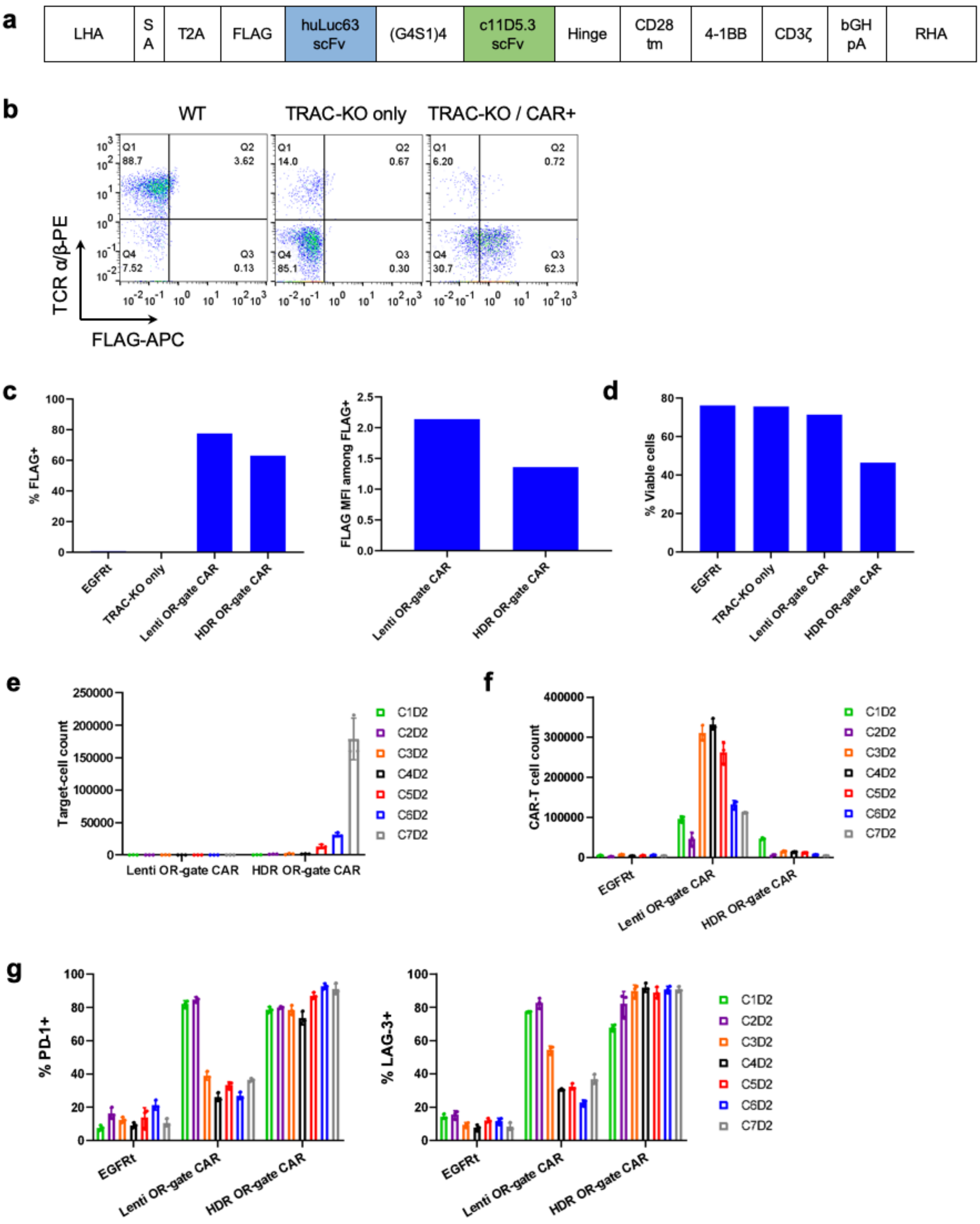


**Supplementary Fig. 17. Lentivirally transduced CAR-T cells outperform HDR-modified CAR-T cells *in vitro*.** **(a)** Schematic of an AAV vector encoding homology arms (left and right, LHA and RHA) flanking an integration cassette consisting of a splice-acceptor (SA) site, a T2A sequence, and the FLAG-tagged huLuc63-c11D5.3 OR-gate CAR. **(b)** Representative TCR α/β and FLAG-tag flow plots 11 days post RNP nucleofection and AAV transduction. **(c)** HDR-modified T cells exhibit lower CAR surface expression than lentivirally-transduced T cells. **(d)** HDR-modified OR-gate CAR-T cells exhibit relatively poor viability 11 days post RNP nucleofection/AAV transduction. **(e–g)** Upon repeated antigen challenge with WT MM.1S cells, HDR-modified OR-gate CAR-T cells exhibit **(e)** inferior cytotoxicity, **(f)** weaker antigen-stimulated T-cell proliferation, and **(g)** stronger and longer-lasting exhaustion-marker (PD-1 and LAG-3) expression than lentivirally transduced OR-gate CAR-T cells. Values shown are the mean of technical triplicate samples with error bars indicating ± 1 SD. Results shown are representative of cells generated from two healthy donors.

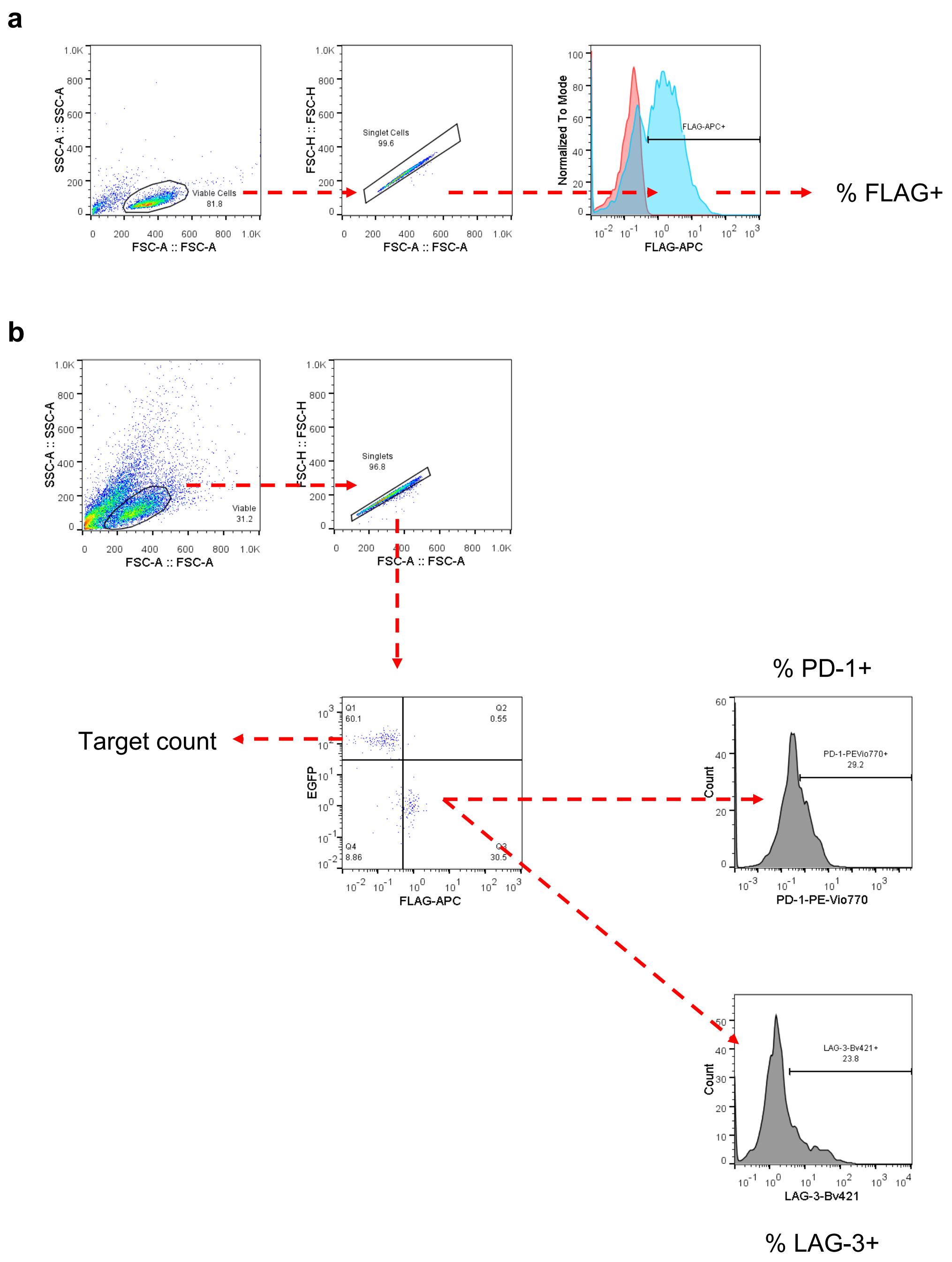


**Supplementary Fig. 18. HDR-modified T cells exhibit poor functional performance in comparison to lentiviral-generated T cells.** Gating strategy for data shown in **(a)** Supplementary Fig. 17c,d, and **(b)** Supplementary Fig. 17e-g.

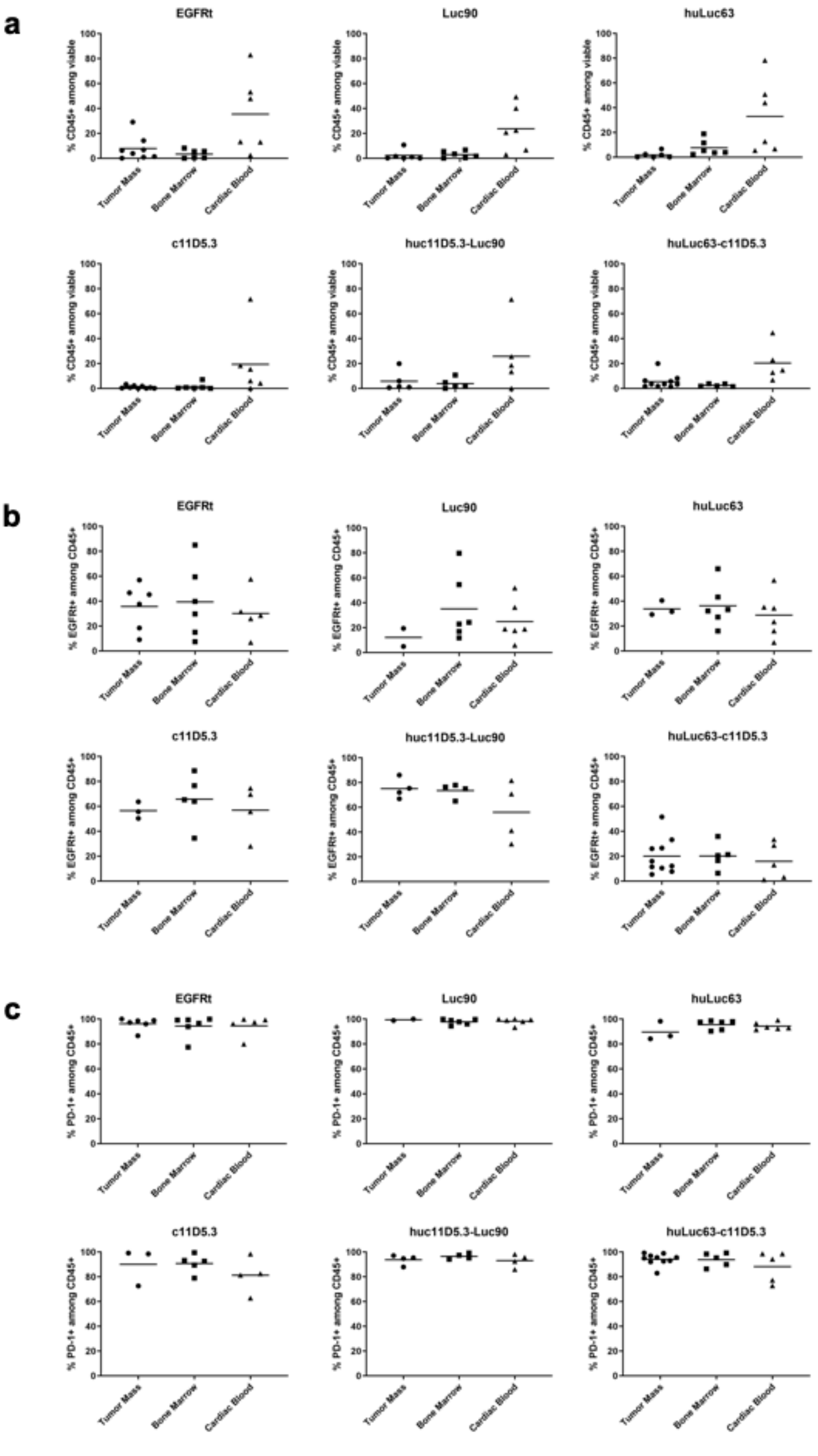
a
EGFRt
Luc90
huLuc63
c11D5.3
huc11D5.3-Luc90
huLuc63-c11D5.3
% CD45+ among viable
Tumor Mass
Bone Marrow
Cardiac Blood
b
% EGFRt+ among CD45+
c
% PD-1+ among CD45+

**Supplementary Fig. 19. CAR-T cells achieve long-term persistent *in vivo* but upregulate PD-1 expression.** Tumor mass, bone marrow, and cardiac blood samples recovered from animals in the study shown in Figure 4 at the time of sacrifice were analyzed for **(a)** the presence of cells expressing human CD45, **(b)** the frequency of $EGFRt^+$ (and thus $CAR^+$) cells among $huCD45^+$ cells, and **(c)** the frequency of PD-$1^+$ cells among $huCD45^+$ cells. Data shown are collected from all the mice in each treatment group (n = 5 or 6), excluding samples that contained <10 $CD45^+$ cells.

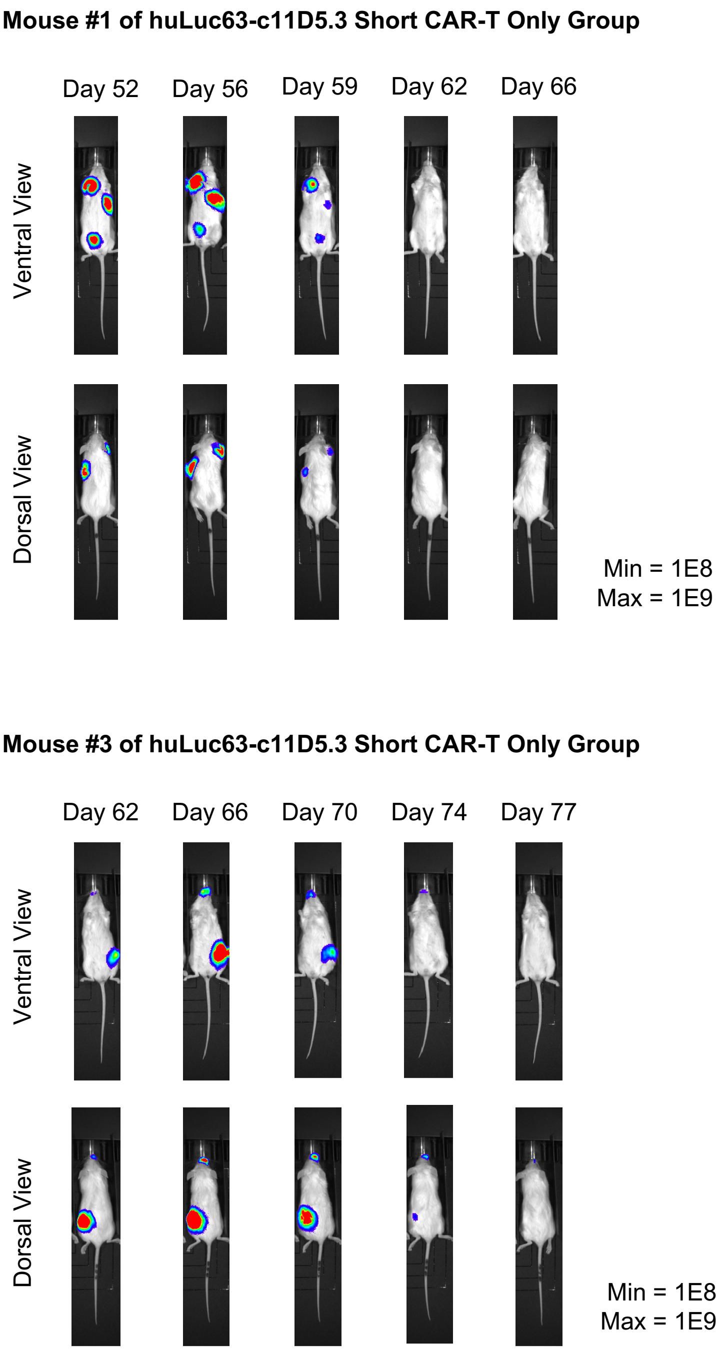


**Supplementary Fig. 20. BCMA/CS1 OR-Gate CAR-T cells can eradicate solid tumor nodules *in vivo*.** NSG mice were engrafted with MM.1S tumors and treated with BCMA/CS1 OR-gate CAR-T cells as described in Figure 6. Two animals in the group treated with huLuc63-c11D5.3 Short CAR-T cells without anti–PD-1 developed palpable solid tumor nodules in the chest and flank, but the tumors eventually regressed to below detection levels and the animals have remained viable in a tumor-free state for >20 weeks, including after tumor re-challenge on day 133, until the study was terminated on day 229. Bioluminescence imaging of the animals in dorsal and ventral views are shown.

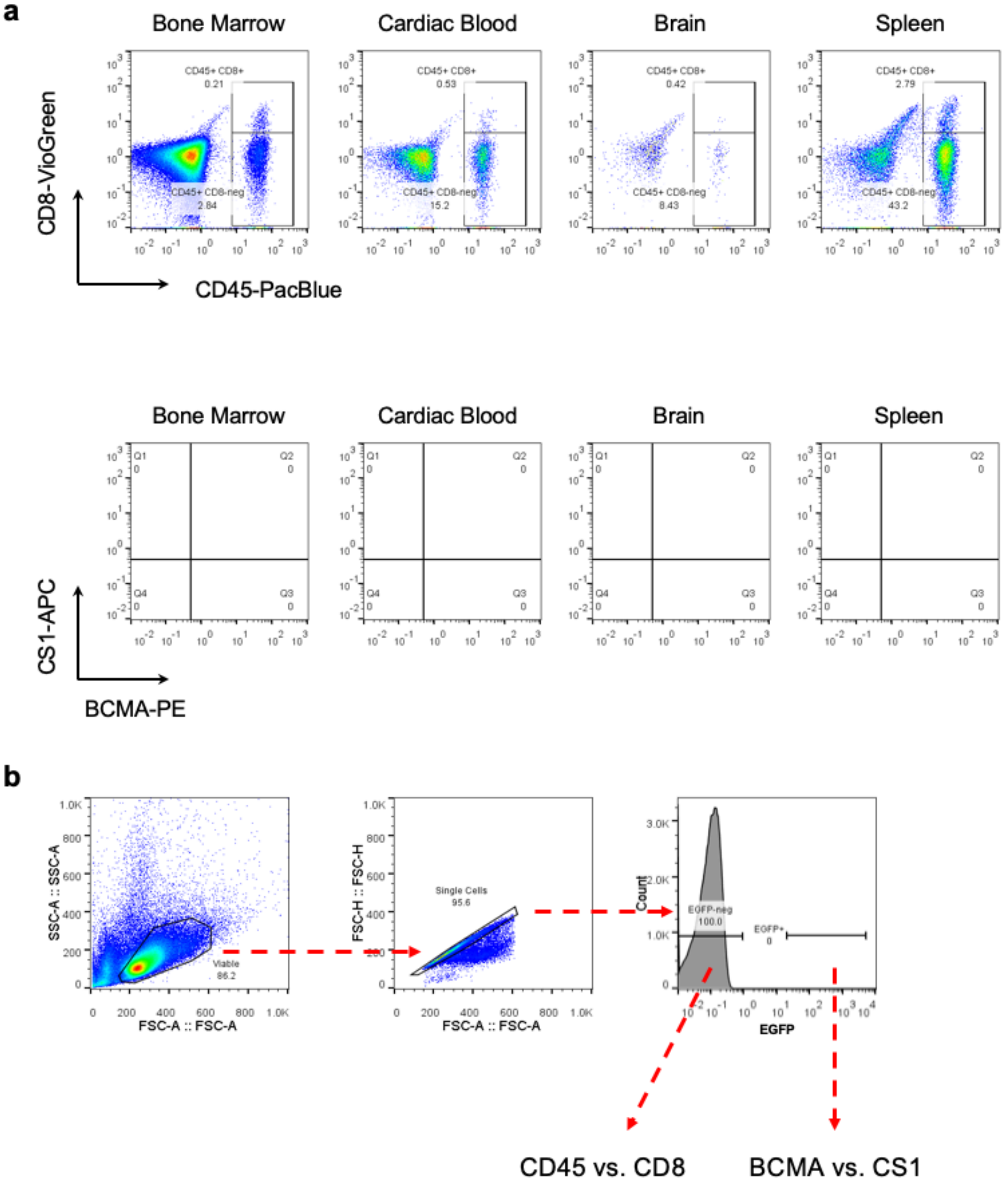


**Supplementary Fig. 21. BCMA/CS1 OR-gate CAR-T cells achieve long-term persistence and prevents tumor relapse *in vivo*. (a)** In the experiment shown in Fig. 6, an animal in the huLuc63-c11D5.3 Short CAR-T cell-treated group without anti–PD-1 was sacrificed on day 165 due to severe weight loss despite the complete absence of tumor signal. This animal had been showing signs of graft-versus-host disease, including fur loss and skin inflammation, for approximately 13 weeks prior to the endpoint. At the time of sacrifice, bone marrow, cardiac blood, brain, and spleen tissue were collected and analyzed by flow cytometry for signs of human T cells and MM.1S tumor cells. Results indicate robust persistence of T cells and the complete absence of tumor cells in this animal. **(b)** Gating strategy for data shown in (a).
